# Mfsd7b facilitates choline uptake and missense mutations affect choline transport function

**DOI:** 10.1101/2023.09.30.560304

**Authors:** Hoa Thi Thuy Ha, Viresh Krishnan Sukumar, Jonathan Wei Bao Chua, Dat T. Nguyen, Toan Q. Nguyen, Lina Hsiu Kim Lim, Amaury Cazenave-Gassiot, Long N. Nguyen

**Affiliations:** Department of Biochemistry, Yong Loo Lin School of Medicine, National University of Singapore, Singapore 119228; Immunology Program, Life Sciences Institute, National University of Singapore, Singapore 117456; Singapore Lipidomics Incubator (SLING), Life Sciences Institute, National University of Singapore, Singapore 117456; Cardiovascular Disease Research (CVD) Programme, Yong Loo Lin School of Medicine, National University of Singapore, Singapore 117545; Immunology Translational Research Program, Yong Loo Lin School of Medicine, National University of Singapore, Singapore 117456

**Keywords:** Choline transporter, Mfsd7b, Flvcr1, PCARP, HSAN, RP

## Abstract

MFSD7b belongs to the Major Facilitator Superfamily of transporters that transport small molecules. Two isoforms of MFSD7b have been identified and they are reported to be heme exporters that play a crucial role in maintaining the cytosolic and mitochondrial heme levels, respectively. Mutations of MFSD7b (also known as FLVCR1) have been linked to retinitis pigmentosa, posterior column ataxia, and hereditary sensory and autonomic neuropathy. Although MFSD7b functions have been linked to heme detoxification by exporting excess heme from erythroid cells, it is ubiquitously expressed with a high level in the kidney, gastrointestinal tract, lungs, liver, and brain. Here, we showed that MFSD7b functions as a facilitative choline transporter. Expression of MFSD7b slightly but significantly increased choline import, while its knockdown reduced choline influx in mammalian cells. The influx of choline transported by MFSD7b is dependent on the expression of choline metabolizing enzymes such as choline kinase (CHKA), but it is independent from gradient of cations. Additionally, we showed that choline transport function of Mfsd7b is conserved from fly to man. Employing our transport assays, we showed that missense mutations of MFSD7b caused reduced choline transport functions. Our results show that MFSD7b functions as a facilitative choline transporter in mammalian cells.

## Introduction

Feline Leukemia Virus Sub-Group C cellular receptor 1 (FLVCR1) was initially discovered as the receptor conferring susceptibility to Feline Leukemia Virus C (FeLV-C) infection in cats and hence named FLVCR1 (1). FLVCR1 also known as MFSD7b belongs to the Major Facilitator Superfamily (MFS) transporters (hereafter referred to as MFSD7b). MFSD7b has twelve transmembrane helices and is ubiquitously expressed (2). Isolated Retinitis Pigmentosa (RP), Posterior Column Ataxia with Retinitis Pigmentosa (PCARP), and Hereditary Sensory and Autonomic Neuropathy (HSAN) have all been linked to the mutations of MFSD7b (3), (4), (5), (6). However, the disease mechanisms are poorly understood and characterized.

Two isoforms of MFSD7b have been identified in human cells which include the smaller 279 amino acid isoform, reported to be localized in the mitochondria (7) and the larger 555 amino acid isoform, which is localized to the plasma membrane (2). Both isoforms are reported to be heme exporters that play a crucial role in maintaining the cytosolic heme pool, particularly during erythropoiesis (8). Studies using *Mfsd7b* knock-out (KO) mice have demonstrated the vital role of MFSD7b in erythropoiesis and how the different isoforms are required for various developmental stages (8), (9), (10). The plasma membrane isoform of Mfsd7b has been shown to be essential for expansion of committed erythroid progenitors (11). The knockout mice of this isoform die at mid-gestation and show disrupted erythropoiesis along with limb and craniofacial deformities which closely resemble patients with Diamond Blackfan Anemia (DBA). Interestingly, mice lacking the plasma membrane isoform, but expressing the mitochondrial isoform of MFSD7b had normal erythropoiesis, implicating that heme export from mitochondria is responsible for the development of blood cells (7). However, the presence of the mitochondrial isoform is not sufficient to prevent hemorhages, edema, and embryonic lethality in mice lacking the plasma membrane isoform. When *Mfsd7b* was knocked out in adult mice, the mice exhibited with severe macrocytic anemia, a phenotype similar to what has been observed in DBA patients (10), (11). Studies performed in zebrafish also show that loss of Mfsd7b disrupts erythropoiesis (11). Global and conditional knockout of *Mfsd7b* in mice exhibit heme accumulation in several cell types like endothelial cells (12), hepatocytes (13), and intestinal cells (14). Morphant knockdown of *mfsd7b* in zebrafish also resulted in heme accumulation (11). These studies support the notion that Mfsd7b has a specific role as a heme exporter that is crucial for survival and proliferation of the cells.

However, the causal link between Mfsd7b role as a heme export and the viability of animals as well as the pathogenesis of patients remains unclear. Heme accumulation is hypothesized to be toxic to the cells. Nevertheless, endogenous heme levels were not increased in MFSD7b mutated patient-derived cells unless when incubated with large amounts of heme or heme precursor 5-aminolevulinic acid. The inconsistency of heme accumulation was linked to the compensatory effects by the increased expression of heme oxidation enzyme (HMOX1). However, in murine erythroid cells, overexpression of Hmox1 to oxidize heme for degradation or reduction of iron to limit heme synthesis in Mfsd7b-knockout cells was not sufficient to rescue proliferative defects due to the loss of Mfsd7b (10). Hence, whether heme accumulation is directly linked to the loss of Mfsd7b function in the human patients and animals remains to be validated.

Given that Mfsd7b is ubiquitously expressed across various organs where its expression is highest in the kidney, gastrointestinal tract, lungs, liver, and brain (8), (7), we hypothesized that Mfsd7b might have additional functions rather than heme export. In this study, we identified additional role of Mfsd7b as a transporter for choline. Consistent with our findings, two recent studies also reported choline transport functions of Mfsd7b (15), (16). Furthermore, we show that Mfsd7b behaves like a facilitative transporter for choline where the expression of choline utilizing enzymes such as choline kinase affects choline import activity. Utilizing this new transport function of Mfsd7b, we show that several missense mutations of MFSD7b found in RP, PCARP, and HSAN patients have a reduction in choline transport activity. Our results implicate that reduced choline transport function in the missense mutations of MFSD7b might be a part of pathogenesis in the patients.

## Results

### Expression of Mfsd7b increases choline uptake

Deletion of Mfsd7b results in heme accumulation (10). Thus, it has been widely accepted that Mfsd7b is a heme exporter. As such, Mfsd7b has been extensively studied for its roles in regulation of erythroid proliferation and differentiation. However, Mfsd7b is ubiquitously expressed in most cell types and tissues (8). Its mRNA levels are higher in the intestine and kidney than the heart and muscle where heme is predicted to be more abundant as a component of the mitochondria in these tissues. In erythroid cells, Mfsd7b is reported to play a role in heme detoxification during differentiation (10). By re-analyzing the gene expression from proerythroblasts to the orthochromatic erythroblasts, we found that expression of Mfsd7b is decreased from the erythroid progenitors to more differentiated cells, while expression of the genes for heme synthesis is increased at later stages of human erythroid cell differentiation (17). Heme levels are also significantly higher in differentiated cells such as orthochromatic erythroblasts than progenitors where Mfsd7c expression is more abundant (10). Notably, we found that expression pattern of Mfsd7b is similar to several metabolic pathways such as mitochondrial activity as well as membrane biosynthesis, but inversely correlated with the heme and hemoglobin synthesis pathway (**Supplemental Figure 1**). Thus, we argue that Mfsd7b might have additional functions rather than being a heme exporter. Interestingly, data from the Cancer Dependency Map project (DepMap) shows that Mfsd7b has a high positive correlation with CHKA and vice versa in CRISPR knockout screens (27). Depmap utilises CRISPR knockout libraries to evaluate the change in fitness across various cancer cell lines for each gene knocked out. A high correlation between two genes represents that loss of either gene has a similar effect on fitness as loss of the other gene, across similar cell lines. This therefore indicates that Mfsd7b and CHKA may have related functions or may play roles in the same pathway. The Kennedy pathway describes the biosynthesis of phosphatidylcholine and phosphatidylethanolamine from choline and ethanolamine (28). CHKA is one of the two choline kinases that phosphorylate choline into phosphorylcholine in this pathway. Thus, we hypothesized that Mfsd7b might transport substrates in the Kennedy pathway. We overexpressed long isoform of human Mfsd7b (hMfsd7b) and its orthologs and performed transport assays in HEK293 cells. Among the tested ligands, we found that hMfsd7b exhibited import activity to choline, but not ethanolamine (**Figure 1A-C**). Choline import activity of hMfsd7b when overexpressed in HEK293 cells was slightly but significantly increased by increasing choline levels in the medium (**Figure 1B**). Mfsd7b is conserved from flies (**Supplemental Figure 2**). We showed that other orthologs of hMFDS7b from fly, fish, chicken, frog, and mouse also exhibited choline transport activity, albeit relatively weak under tested conditions (**Figure 1A**). hMfsd7b did not transport L-carnitine, acetylcholine, betaine, and serotonin (**Figure 1C**). These results hint that Mfsd7b might play a role in choline transport. Two recent reports also show that FLVCR1/Mfsd7b exhibits choline transport activity which is in line with our observations (15), (16).

**Figure 1.**
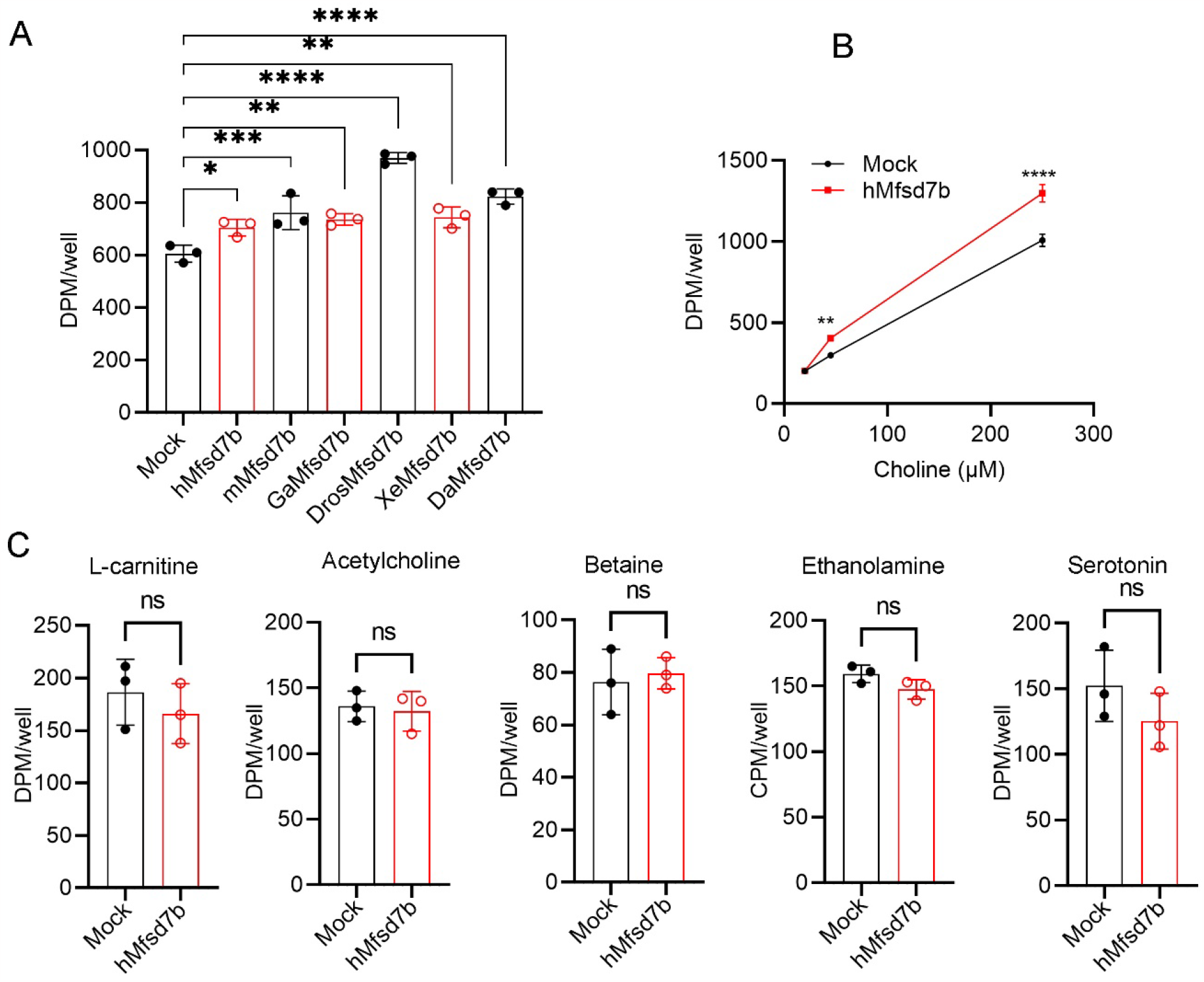
Expression of Mfsd7b increases choline import. **A**, overexpression of Mfsd7b increases choline uptake. HEK293 cells were overexpressed with human (hMfsd7b), mouse (mMfsd7b), chicken (GaMfsd7b), fly (DrosMfsd7b), frog (XeMfsd7b), and fish (DaMfsd7b) Mfsd7b orthologs and tested for choline uptake assay. Mock was transfected with empty plasmid. **B**, Dose curve of choline import by hMfsd7b in HEK293 cells. **C**, Import assays for hMfsd7b with indicated ligands. Experiments were performed at least twice in triplicate. Data are mean and SD. ****P<0.0001, ***P<0.001, **P<0.01, *P<0.05; One-way ANOVA for A and Two-way ANOVA for B; ns, not significant.

### Deficiency of Mfsd7b results in reduced choline uptake

To further demonstrate that Mfsd7b regulates choline influx, we knocked down Mfsd7b transcripts using siRNA to avoid potential cell viability phenotype that has been reported in complete knockout of Mfsd7b previously (18) and used the knockdown cells for transport assays. We chose A549 cell line as Mfsd7b mRNA is expressed in this cell type (2). We first confirmed that siRNA could sufficiently knock down Mfsd7b transcripts to decrease Mfsd7b protein levels in A549 cells (**Figure 2A**). Then, we performed choline transport assays of the wildtype and knockdown cells. Our results showed that A549 cells with deficiency in Mfsd7b exhibited significantly reduced choline import compared to controls (**Figure 2B**). Additionally, we showed a reduction of choline uptake which resulted in slightly but significant reduction of phosphatidylcholine (PC) and sphingomyelin (SM) levels in Mfsd7b knockdown compared to control A549 cells (**Figure 2C, D; Supplemental information table 1**). These results implicate that choline imported by Mfsd7b may be used to phospholipid synthesis in the cell line.

**Figure 2.**
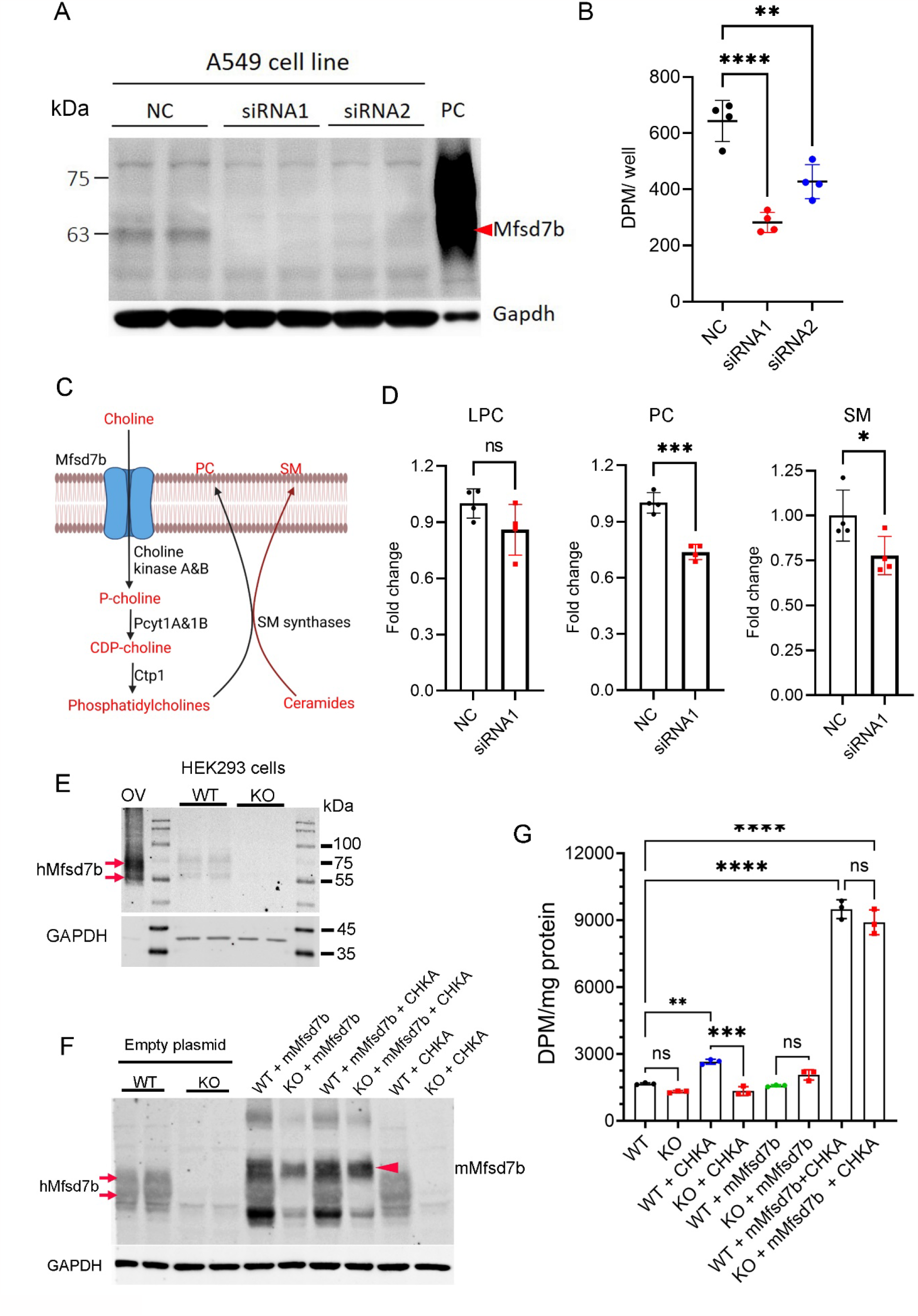
Deletion of Mfsd7b reduces choline transport in mammalian cells. **A**, Western blot analysis of siRNA knockdown of Mfsd7b in human cell line A549. Experiments were repeated twice with duplicate. Red arrowhead indicates Mfsd7b band. PC, positive control from overexpression of hMfsd7b in HEK293 cells. NC, siRNA negative control. **B**, Import of choline is significantly reduced in A549 cells after Mfsd7b is knocked down by siRNA. Experiments were repeated at least twice in 4 replicates. **C**, Illustration of import of choline by Mfsd7b and conversion of imported choline to phosphatidylcholine via Kennedy pathway. **D**, Lipidomics analysis shows Phosphatidylcholine level is significantly reduced in A549 cells after Mfsd7b is knocked down by siRNA. **E**, generation of Mfsd7b knockout in HEK293 cells using CRISPR/Cas9 method. Expression of Mfsd7b is abolished in the knockout cells (KO). (OV: overexpression of Mfsd7b used as positive control), **F**, Rescue experiments in Mfsd7b-KO HEK293 cells. Wildtype and KO cells were expressed with mouse Mfsd7b (mMfsd7b) or co-expressed with mouse Mfsd7b and CHKA. **G**, Choline transport activity of WT and KO cells with expression of mMfsd7b, CHKA or co-expression of Mfsd7b with CHKA as shown in G. ****P<0.0001, **P<0.01; *P<0.05. One-way ANOVA in B; t-test in D.

Recently, Kenny et al. reported that overexpression of Mfsd7b in Mfsd7b-knockout cells strongly drives choline import in HEK293 cells, whereas Tsuchiya et al. performed similar experiments but did not observe the increased choline transport activity beyond the wild-type levels (16), (15). Since we showed that overexpression of hMfsd7b in wild-type cells only modestly increases choline import activity, we wanted to validate our results using Mfsd7b-knockout cells. To this end, we generated Mfsd7b knockout in HEK293 cells using CRISPR/Cas9 technology and employed the cells for our rescue experiments (**Figure 2E**). Consistent with the results using wild-type HEK293 cells, we showed that overexpression of mouse Mfsd7b (mMfsd7b) in these KO HEK293 cells did not significantly enhance choline import (**Figure 2F-G**). Interestingly, co-expression of mMfsd7b with CHKA in wild-type and KO HEK293 cells significantly increased choline uptake (**Figure 2F-G**). These results strongly indicate that Mfsd7b facilitates choline uptake in cells, and that choline influx mediated by Mfsd7b is likely governed by intracellular choline utilizing enzymes.

### Choline imported by Mfsd7b is increased by the expression of choline metabolizing enzymes

We observed that although heterologous expression of Mfsd7b in HEK293 cells increased choline uptake, it did not greatly increase choline influx at higher doses (**Figure 1B**). These results suggest that the rate of utilizing intracellular choline is perhaps a limiting factor that affects choline transport by Mfsd7b. Indeed, co-expression of Mfsd7b with choline kinase A (CHKA) to convert choline into phosphorylcholine greatly enhanced choline uptake (**Figure 2F-G**). To test if lowering the intracellular levels of choline is sufficient to increase choline import mediated by Mfsd7b, we also used choline acetyltransferase (CHAT) to facilitate with the conversion of choline into acetylcholine. By lowering the intracellular levels of choline to phosphorylcholine with the co-expression of CHKA or acetylcholine with the co-expression of CHAT, we showed that the influx of choline was significantly increased (**Figure 3A-B**). Co-expression of CHKA with Mfsd7b orthologs from fly, frog, fish, chicken, and mouse also significantly increased import of choline in a similar capacity to human Mfsd7b (**Figure 3C**). Furthermore, choline uptake by Mfsd7b was greatly enhanced with the increased concentrations of choline when co-expressed with CHKA (**Figure 3D**). The import of choline by hMfsd7b was also significantly increased over time (**Figure 3E**). We also included the N121D mutant of Mfsd7b from a PCARP patient and showed that choline uptake activity of the mutant was significantly reduced compared to that of WT protein (**Figure 3D-E**). Nevertheless, Mfsd7b did not import ethanolamine even with the co-expression of ETNK1, suggesting that it might have specificity to choline (**Figure 3F**). To gain further insight into the new role of Mfsd7b as a choline transporter, we performed complementation experiments. We showed that expression of zebrafish Mfsd7b or CHT1 rescued the reduction of choline uptake when human Mfsd7b is knocked down in HEK293 cells (**Supplemental Figure 3A**). Additionally, expression of hMfsd7b rescued for the reduction of choline transport mediated by neuronal choline transporter CHT1 in HEK293 cells (**Supplemental Figure 3B**).

**Figure 3.**
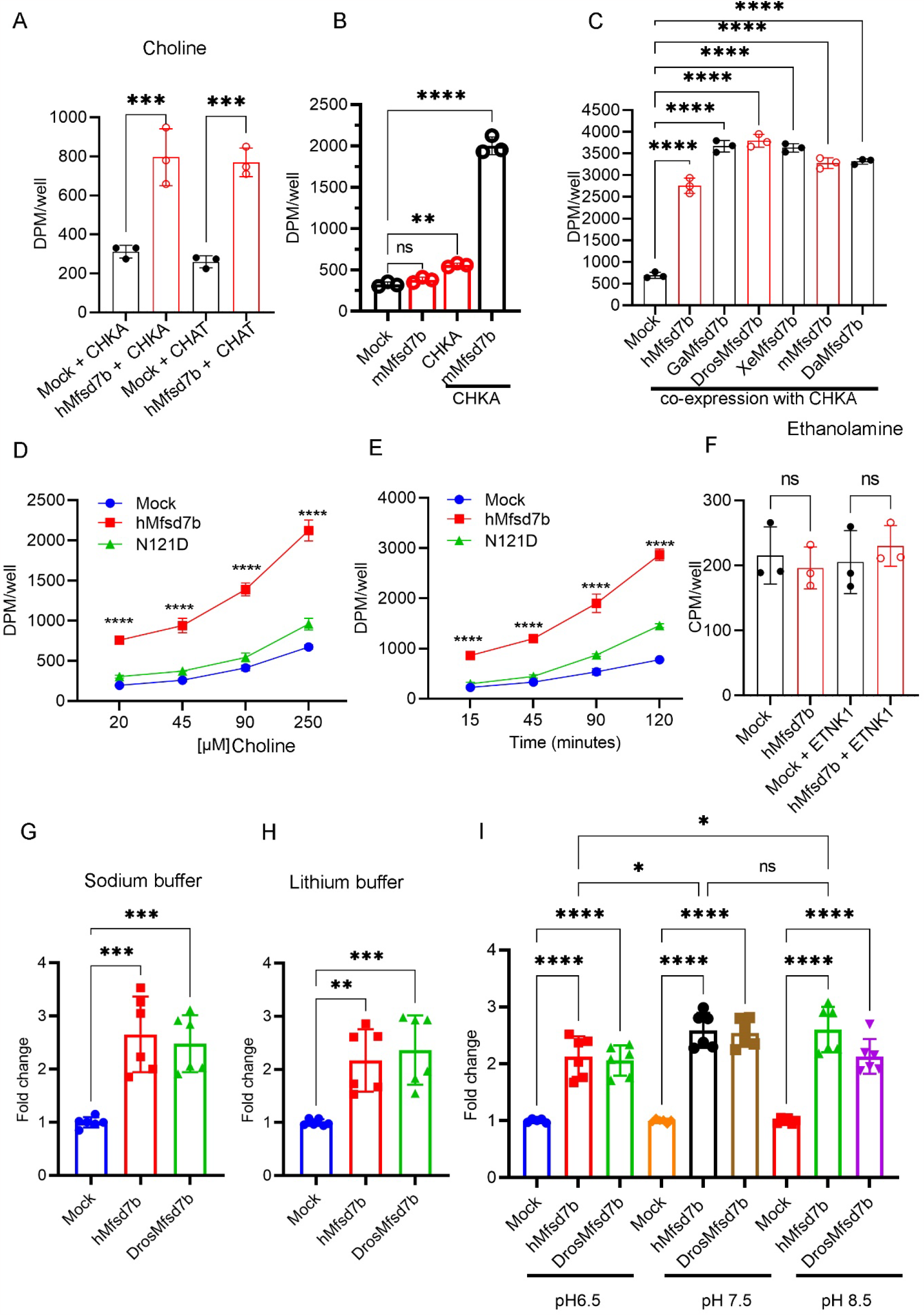
Expression of choline metabolizing enzymes increases choline influx via Mfsd7b. **A**, Co-expression of choline metabolizing enzymes, CHKA and CHAT significantly increases choline uptake via Mfsd7b. HEK293 cells were transfected with hMfsd7b and CHKA or hMfsd7b and CHAT. Mock was transfected with CHKA or CHAT alone. **B-C**, Co-expression of CHKA with Mfsd7b orthologs greatly increases choline uptake. **D**, Co-expression of hMfsd7b with ethanolamine kinase 1 (ETNK1) does not increase uptake of ethanolamine. **E**, Transport activity of hMfsd7b with different doses of choline is significantly increased when co-expressed with CHKA. **F**, Uptake of choline by hMfsd7b is increased over time when co-expressed with CHKA. **G-H**, Choline import activity of hMfsd7b and DrosMfsd7b is not sodium dependent. Choline uptake by hMfsd7b and DrosMfsd7b relative to mock is not affected by presence or absence of sodium. **I**, Choline uptake by hMfsd7b and DrosMfsd7b is modestly affected by pH. Data are mean and SD. ****P<0.0001, ***P<0.001; One-way ANOVA for A-C and F-I and Two-way ANOVA for D-E; ns, not significant.

Next, we tested whether Mfsd7b requires a gradient of cations for transport of choline. Replacement of sodium with lithium or changing pH in the transport buffer did not affect choline import activity of Mfsd7b (**Figure 3G-I**). These results demonstrate that Mfsd7b is a transporter for choline and the influx of choline facilitated by the expression of choline metabolizing enzymes is not dependent on the gradient of cations. Collectively, our results show that Mfsd7b is a conserved choline transporter that facilitates choline transport via the plasma membrane.

### Missense mutations of Mfsd7b cause reduced choline uptake activity

Several missense mutations of Mfsd7b have been reported in patients with retinitis pigmentosa (RP) and posterior column ataxia with retinitis pigmentosa (PCARP), and hereditary sensory and autonomic neuropathy (HSAN). However, it is unclear if choline transport function of Mfsd7b is affected by these mutations. Thus, we generated these missense mutations by mutagenesis and tested their transport activity (**Supplemental Table 1**, list of mutants). We found that all of these missense mutations of Mfsd7b resulted in reduced choline import activity (**Figure 4A**). Several missense mutants such as N121D and L160P had abolished or severely reduced choline transport activity (**Figure 4A**). In comparison with the transport activity of native protein, these mutants retained 0-57% transport activity (**Figure 4B**). Mfsd7b has two potential N-glycosylation sites (N265 and N273). We mutated these residues to alanine (N265A and N273A) and tested their transport activity. We found that these mutant proteins hade slightly reduced choline transport activity (**Figure 4C**). The reduced choline transport activity of these mutated proteins was not due to reduced expression levels (**Figure 4D**) and localization (except for C192R, L199P, A241T, A283P, and Y341C mutants) (**Supplemental Figure 4**). These data show that the change of amino acids due to the missense mutations in Mfsd7b causes reduced choline transport activity. If the transport activity for both alleles of Mfsd7b is calculated, the net of transport activity of Mfsd7b mutants is lower than 41% in these patients, except for patient 4 and 8 with compound heterozygous mutations (**Supplemental Table 1**). These results suggest that reduced choline transport activity in these mutations of Mfsd7b may be a part of the pathogenesis of the patients.

**Figure 4.**
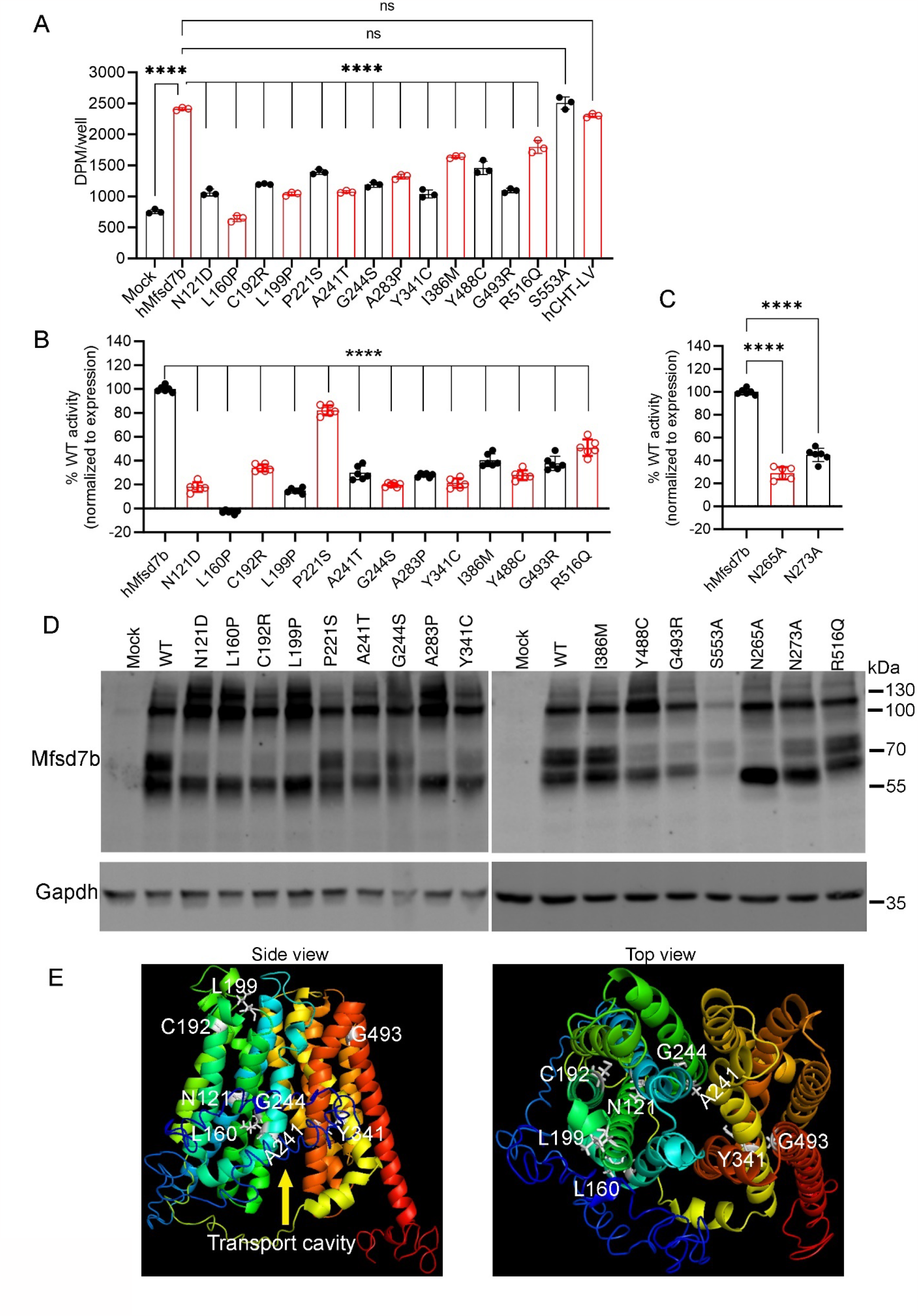
Missense mutations of Mfsd7b found in RP, PCARP, HSAN patients cause reduced choline transport. **A**, transport activity of indicated missense mutations of hMfsd7b with choline. The missense mutants and native hMfsd7b were co-expressed with CHKA in HEK293 cells. hCHT-LV is the human neuronal choline transporter (also called SLC5a7) dominant active mutant (with dileucine motif LV mutated to AA) was used for positive control. Mock was transfected with CHKA alone. **B**, Transport activity of indicated missense mutants expressed as percentage of wild-type hMfsd7b (WT) activity. Transport activity of the mutants was normalized to the protein expression quantified from western blots. **C**, Transport activity of two mutants for glycosylated sites N265A and N273A. Experiments were performed at least twice in triplicate. **D**, Western blot analysis of mutated proteins from overexpression in HEK293 cells. WT denotes native hMfsd7b. Data are mean and SD. ****P<0.0001, ***P<0.001; One-way ANOVA for A-C. ns, not significant. **E**, Side and top views of a structural model of hMFSD7b. Amino acids shown in grey are present in the missense mutations which resulted in the choline transport activity below 30% of that of WT protein. Except for L160P and G244S mutants, the remaining mutants came from patients with homozygous missense mutations of MFSD7b. Arrow indicates predicted transport cavity of hMFSD7b.

## Discussion

FLVCR1/MFSD7b was discovered as a receptor for the envelope protein of Feline Leukemia Virus Sub-Group C that causes leukemia in cats. Its molecular functions were then revealed as a heme exporter. Heme detoxification by Mfsd7b is shown to be important for cell viability. This process is thought to be mainly active in erythroid lineage because these blood cells employ Mfsd7b to maintain a balance level of heme for hemoglobin synthesis during erythropoiesis. Indeed, erythroid cells lacking Mfsd7b ceased proliferation that results in severe anemia in the knockout mice (9), (10). However, Mfsd7b is ubiquitously expressed in different cell types in mice (8), (7). Mfsd7b is also a conserved protein found in flies which do not utilize hemoglobin for the transport of oxygen. These lines of evidence suggest that Mfsd7b might have additional molecular roles along with heme export function. Our results here show that the 555 amino acid long, plasma membrane isoform of human Mfsd7b is a transporter for choline.

Employing gene deletion and overexpression in mammalian cells, we show that Mfsd7b exerts choline transport function, but not other structurally similar molecules such as betaine and acetylcholine. Our results also show that heterologous expression of several Mfsd7b orthologs from flies to mouse in mammalian cells also increases the import of choline. Thus, choline transport function of the long isoform Mfsd7b is conserved. Interestingly, choline uptake by Mfsd7b is significantly enhanced when choline kinase A (CHKA) or choline acetyltransferase (CHAT) is co-expressed, indicating that increasing the levels of intracellular choline affects the influx of choline facilitated by Mfsd7b. Our results show that a gradient of cations such as sodium and proton is not required for choline transport activity of Mfsd7c. These results strongly suggest that Mfsd7b facilitates the uptake of choline downhill of its concentration. The influx of choline mediated by Mfsd7b is also regulated by intracellular choline utilizing enzymes.

Whole body deletion of Mfsd7b in mice results in embryonic lethality (7), (8). Additionally, conditional deletion of Mfsd7b using Mx1Cre also results in lethality due to severe anemia (10). Patients with mutations of Mfsd7b have been reported to have eye disease (retinitis pigmentosa), degeneration of spinal cord (posterior column ataxia), and compromised sensory system (hereditary sensory and autonomic neuropathy). However, anemic symptoms were not reported in these patients. Using choline transport assay, our results show that most of the missense mutations of the long isoform of hMfsd7b results in significant reduction of choline transport activity. Several missense mutations such as N121D, L160P, and L199P have less than 20% choline transport activity compared to the native protein. It is also noteworthy that several inactive mutations such as N121D are only present in the long isoform of hMfsd7b. Thus, it might rule out the active role for choline transport function of the short isoform. Hence, these results suggest that reduced choline transport activity due to the mutations of hMFSD7b may be a part of the pathogenesis in the patients. Nevertheless, these findings will need validations in vivo settings.

An important question is that whether the reduction or loss of heme export function of Mfsd7b is the causal factor for the symptoms in the human patients. To our knowledge, heme export activity has not been tested for the mutated proteins in the patients. We argue that if heme accumulation is the only pathological factor, the patients with *HMOX1* mutations would also exhibit similar symptoms. However, patients with HMOX1 mutations exhibit increased oxidative stress (19), hematological symptoms and inflammation (20), and lung disease (21). Additionally, global deletion of Hmox1 in mice do not affect viability (22). In support of our notion for the involvement of choline as one of the causal factors, human patients with mutations of MFSD7b appear to have overlapping symptoms with the patients who carry mutations in the genes of the Kennedy pathway (23). For example, mutations of Pcyt1a which is the rate-limiting enzyme for phospholipid synthesis from choline in the Kennedy pathway causes eye disease (23). Therefore, we speculate that reduced choline transport function caused by mutations of MFSD7b could be a part of the pathogenesis in the patients. We believe that the identification of MFSD7b as a choline transporter is an important step for future research to reveal the disease mechanisms and possible treatments for the patients.

## Experimental procedures

### Cell culture

Human embryonic kidney (HEK293) and A549 cell lines were maintained in Dulbecco’s Modified Eagle Medium (Thermo Fisher Scientific, Lithuania) containing 10% heat-inactivated fetal bovine serum (Thermo Fisher Scientific), 50 μg/mL penicillin (Thermo Fisher Scientific), and 50 μg/mL streptomycin (Thermo Fisher Scientific). Cells were maintained in an incubator at 37°C and 5% CO_2_.

### Plasmids and mutagenesis

Human MFSD7b plasmid was described previously (8). cDNA of long isoform of human MFSD7b *(hMFSD7b*, NM_014053.4) was cloned into pcDNA3.1 for overexpression. Mouse *(mMfsd7b*, NM_001081259.1), chicken *(GaMfsd7b*, XM_419425.7), fly *(DrosMfsd7b*, NM_001201944.2), frog *(XeMfsd7b*, XM_002934686.5), and fish *(DaMfsd7b*, NM_001025522.1*) Mfsd7b* coding sequences were synthesized by GeneScript and cloned into pcDNA3.1 plasmid. To generate the missense mutations, hMFSD7b cDNA was used as a template for mutagenesis by PCR. The cDNA carrying each missense mutation was cloned into pcDNA3.1 and confirmed by Sanger sequencing.

### Antibodies

In-house polyclonal antibodies against human MFSD7b were raised in rabbits using 13 amino acids KMVMLSKQSESAI at the C-terminus of human MFSD7b protein. Other antibodies used were commercially available.

### Transport assays in HEK293 cells

To test the transport activity of hMFSD7b or Mfsd7b orthologs, the cDNA of Mfsd7b gene in pcDNA3.1 was transfected to HEK293 cells using lipofectamine 2000. Mock was transfected with empty plasmid. For experiments in which human choline kinase A (CHKA) was used, 1.5μg pcDNA3.1 CHKA plasmid was co-transfected with 2.5μg plasmid of hMfsd7b or Mfsd7b orthologs or mutant plasmids. Cells transfected with 1.5μg pcDNA3.1 CHKA plasmid was used as control. After 20-30 hours post-transfection, HEK293 cells were used for transport assays with 100μM [^3^H] choline in DMEM containing 10%FBS. The transport assays were stopped after 1hour of incubation at 37°C by washing once with cold plain DMEM medium. Cell pellets were lyzed in RIPA buffer and transferred to scintillation vials for quantification of radioactive signal using Tricarb liquid scintillation counter. Transport activity of Mfsd7b was expressed as DMP for [^3^H] isotopes and CPM for [^14^C] isotopes. For dose-dependent transport activity, HEK293 cells were similarly co-transfected with wildtype hMfsd7b, or hMfsd7bN121D mutant plasmid with CHKA plasmid. Transport activity was conducted with 20, 45, 90, and 250 μM [^3^H] choline in DMEM containing 10% FBS for 1 hour at 37^0^C. For time-dependent assay, hMfsd7b or hMfsd7bN121D was co-transfected with CHKA. Transfected HEK293 cells were assayed with 100μM [^3^H] choline in DMEM containing 10%FBS for 15, 30, 60, and 120 mins at 37^0^C. For transport assays with [^14^C] ethanolamine, [^3^H] L-carnitine, and [^3^H] betaine, [^3^H] acetylcholine, and [^3^H] serotonin, HEK293 cells were co-transfected with hMfsd7b and CHKA as described above. Transport assays were performed with 100μM [^14^C] ethanolamine, 100μM [^3^H] betaine, [^3^H] acetylcholine, [^3^H] serotonin, and 50 μM [^3^H] L-carnitine, in DMEM with 10%FBS for 1 hour. Cells were washed once with cold plain DMEM and lyzed with RIPA buffer for radioactive quantification. Choline import assays for A549 knockdown cell line were similarly performed. Briefly, after 48h of siRNA transfection, Mfsd7b-knockdown cells and controls were incubated with 100 μM [^3^H] choline for 2 hours. Cells were washed once with cold plain DMEM and lyzed with RIPA buffer for scintillation quantification. For sodium and pH-dependent experiments, the buffer containing 0.54 mM KCl, 1.3 mM CaCl2, 0.53 mM MgCl2, 0.4 mM MgSO4, 0.37 mM KH2PO4, 138 mM NaCl, 0.28 mM Na2HPO4, 5.5 mM Glycine, 5.6 mM D-Glucose at pH 6.5, 7.4 or 8.5 was used (24). For lithium containing buffer, sodium salts in the transport buffer were replaced with 138mM of LiCl.

### Western blot

Overnight transfected HEK293 cells were lysed in RIPA buffer (25 mM Tris pH 7–8, 150 mM NaCl, 0.1% SDS, 0.5% sodium deoxycholate 0.5% Triton X-100) with protease inhibitor (Roche, 11836170001). Protein quantification was done by Pierce BCA protein assay following manufacturer’s protocol. After total protein quantification, 10 μg of total protein lysate was resolved in 10% SDS-PAGE and transferred to 0.45 μm nitrocellulose membrane. The membranes were probed with rabbit anti-human MFSD7b primary antibody and IRDye® 800CW Donkey anti-Rabbit IgG (LI-COR Biosciences, 926-32213) secondary antibody. The membranes were reprobed with mouse anti-GAPDH primary antibody (Santa Cruz, sc-32233) and IRDye® 680LT Goat (polyclonal) anti-Mouse IgG (H+L) (LI-COR Biosciences, 926-68020) secondary antibody. PageRuler Plus Prestained Protein Ladder (Thermo Scientific, 26620) was used as protein size standard. The membranes were imaged by ChemiDoc MP Imaging system (Bio-Rad).

### Immunofluorescence microscopy

HEK293 cells were seeded on glass coverslips in 12 well plates and transfected with hMFSD7b or mutant plasmids with the plasma membrane localized GFP (mGFP) plasmids. At 24 hours post transfection, the transfected cells were fixed in 4% paraformaldehyde for 30 minutes. The fixed cells were then washed with PBS and permeabilised in 0.1% Triton-X in PBS for 15 minutes. Next, the cells were incubated with 5% Natural Goat Serum (NGS) in 0.1% Triton-X in PBS for 15 minutes. The cells were then incubated with primary antibody rabbit anti-human MFSD7b in 5% NGS in 0.1% Triton-X in PBS for 1 hour. The cells were washed with PBS and incubated with secondary antibody Goat Anti-Rabbit IgG (H+L), Alexa Fluor® 488 conjugate Antibody (Life technologies, A11034) in 5% NGS in 0.1% Triton-X in PBS for 1 hour. Afterwards, the cells were washed in PBS and incubated in Hoechst 33342 in PBS for 5 minutes then washed again with PBS. The glass coverslips were transferred to poly-lysine coated slides and covered in mounting media (70% glycerol, 20mM Tris HCL, pH 6). The slides were covered with cover glass and sealed with nail polish. The slides were visualized by Zeiss LSM 710 Confocal Microscope at 63x magnification.

### Knockdown of MFSD7b with siRNA in A549 and HEK293 cells and knockout of MFSD7b in HEK293 cells

Double stranded siRNAs targeting hMFSD7b were purchased from IDT (Integrated DNA technologies, Singapore). A549 cells were seeded and transfected with siRNA (for MFSD7b or negative control) using transfection reagent Lipofectamine RNAiMAX (Invitrogen) following the manufacturer protocol. After 48 hours of transfection, cells were used for transport assays and Western blot analysis. For HEK293 cells, the cells were seeded and transfected with siRNA using transfection reagent Lipofectamine RNAiMAX (Invitrogen) following the manufacturer protocol. After 24 hours of siRNA transfection, the cells were transfected with pcDNA3.1 vector containing zebrafish MFSD7b (DaMfsd7b) or CHT1-LV mutant gene to rescue the knockdown of hMFSD7b. After 4 hours of the plasmid transfections, the cells were transfected with another dose of siRNAs to enhance gene knockdown efficiency. After 48 hours of the first siRNA transfection, the cells were used for transport assays. For knockout of MFSD7b in HEK293 cells, we employed CRISPR/Cas9 technology. The absence of Mfsd7b protein was validated by Western blot.

### Lipidomic analysis

Wildtype and siRNA1 knockdown A549 cells were grown in 12-well plates and then harvested using cell scrappers in 1 ml PBS/well. The cells were then centrifuged at 3,000 ×g for 5 minutes to collect the cell pellets. An amount of 200 μl butanol/methanol (1:1, v/v) containing internal standards (Avanti Lipids) was added in each sample to extract lipids. The samples were then sonicated in a water bath for 30 minutes, followed by centrifugation at 14,000 ×g for 10 minutes. The extracted lipids were then separated using a Hilic column (Kinetex 2.6 μm HILIC 100 Å, 150 × 2.1 mm, Phenomenex, Torrance, CA, USA) on an Agilent 6495A2 (Agilent Technologies). Mobile phases A (50% acetonitrile (LC-MS grade, Thermo Fisher Scientific Inc.) 50% 25 mM ammonium formate (Sigma-Aldrich) pH = 4.6) and B (95% acetonitrile and 5% 25 mM ammonium formate pH = 4.6) were mixed at the following gradient: 0–6 minutes, 99–75% B; 6–7 minutes, 75-10% B; 7–7.1 minutes, 10–99.9% B; 7.1–10.1 minutes, 99.9% B. The raw data were processed using Agilent MassHunter Quantitative Analysis software for QQQ version B.08 (Agilent Technologies, Santa Clara, CA, USA). The lipid samples were normalized to relevant internal standards and protein concentration. Mass spectrometer parameters were as follows: electrospray ionization, gas temperature 300°C, gas flow 5 L/minute, sheath gas flow 11 L/minute, and capillary voltage 3,500 V. Phospholipids were quantified at the sum composition level using multiple reaction monitoring (MRM), see supplementary information table 1 for detailed list of MRMs.

### Transport assay in HEK293 cells with the presence of hemicholinium-3 (HC-3)

HEK 293 cells transfected with human CHKA (Mock), human CHKA + human MFSD7B, human CHKA + human CHT1-LV, or human CHKA + fly Mfsd7b (DrosMfsd7b) were incubated with 100 μM [3H] choline or 100 μM [3H] choline + 200 μM hemicholinium-3 in DMEM containing 10% FBS at 37°C for 1 hour. Subsequently, the assay was stopped by removing the medium containing [3H] choline, and washed once with cold plain DMEM. The cells were then lyzed in RIPA buffer for 15 mins, and transferred to scintillation vials for radioactive quantification using Tricarb liquid scintillation counter.

### Statistical Analyses

Results were analysed using GraphPad Prism, version 7.2, software for Windows. Statistical significance was calculated using one-way ANOVA test, as indicated in the figure legends. A *P* value of less than 0.05 was considered statistically significant.

## Data availability

All data are available upon request. All data have been included in the article.

## Acknowledgements

We thank Dr. Raymond Doty for the plasmid gift.

## Author contributions

T.Q.N., T.D.N., J.C., and L.N.N. performed *in vitro* transport assays. H.T.T.H. and J.C.W.B. performed siRNA knockdown and knockout experiments and transport assays. V.K.S. generated the mutant plasmids and WB results. H.T.T.H. performed IF experiments. A.C-G. performed lipidomics. L.L. provided reagents. L.N.N. conceived and designed the study and experiments; analyzed the data, prepared the figures, and wrote the paper with V.K.S.

## Supporting information

**Supplemental Figure 1.**
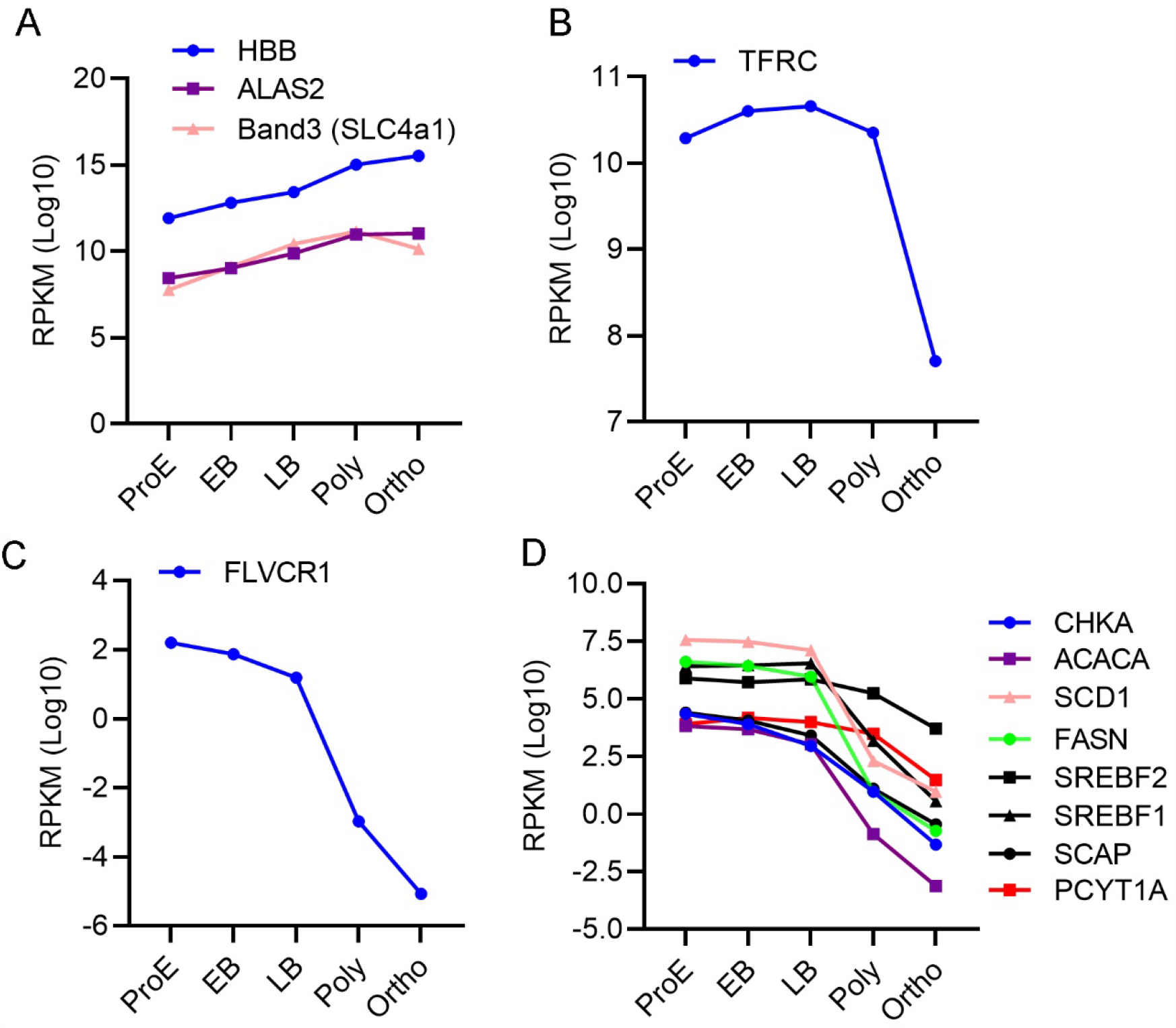
Expression of FLVCR1/Mfsd7b during erythropoiesis. **A**, increased expression of heme and hemoglobin synthesis gene HBB and ALAS2, respectively. Expression of Band3 gene is also increased as it is required to maintain the levels of anions in the red blood cells. **B**, Expression of the transferrin receptor TFRC which is required for iron delivery to erythroid cells. **C**, Expression of FLVCR1 is decreased during erythropoiesis. The expression pattern of FLVCR1 (Mfsd7b) is inversely correlated with the expression of heme and hemoglobin synthesis enzymes shown in A. **D**, Expression of choline kinase A (CHKA), acetyl-CoA carboxylase (ACACA), fatty acid desaturase 1 (SCD1), fatty acid synthase (FASN), sterol responsive element binding protein 1 and 2 (SREBF1 and 2), SREBPF-1/2 binding protein (SCAP), and choline-phosphate cytidylyltransferase A (PCYT1A) is dereased during erythropoiesis. Cellular stages during erythropoiesis: Proerythroblast (ProE), Early and late basophilic erythroblast (EB and LB), Polychromatic erythroblast (Poly), Orthochomatic erythroblast (Ortho). The above data were derived from Xiuli An et. al., 2014.

**Supplemental Figure 2.**
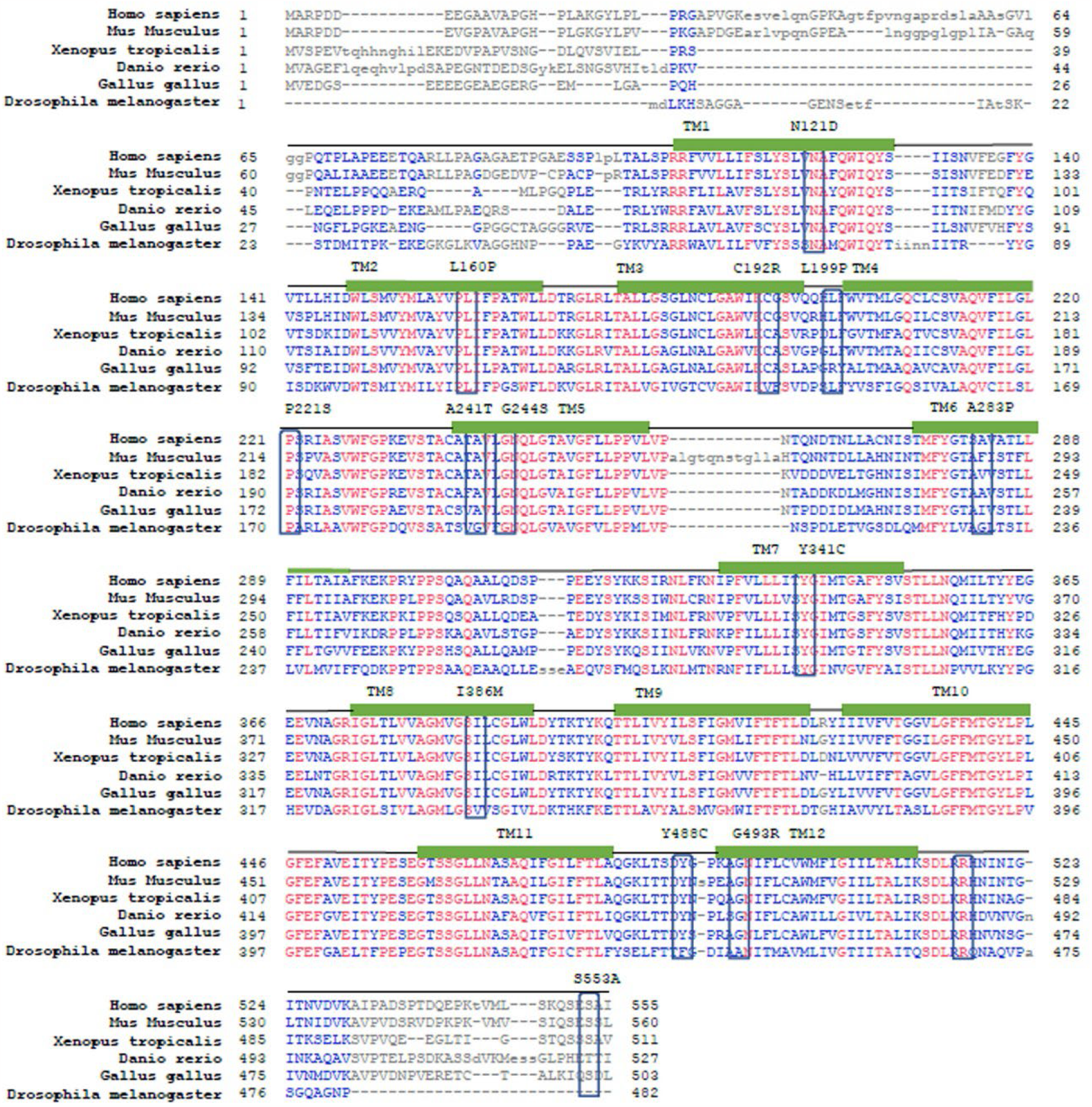
Amino acid sequence alignment of human, mouse, frog, zebrafish, chicken and fly Mfsd7b. Mfsd7b is a conserved protein from flies which has 12 predicated transmembrane domains (TM1-12). Several missense mutations of Mfsd7b have been identified which cause Retinitis Pigmentosa (RP), Posterior Column Ataxia with Retinitis Pigmentosa (PCARP), and Hereditary Sensory and Autonomic Neuropathy (HSAN). Most of these missense mutations are conserved across the species.

**Supplemental Figure 3.**
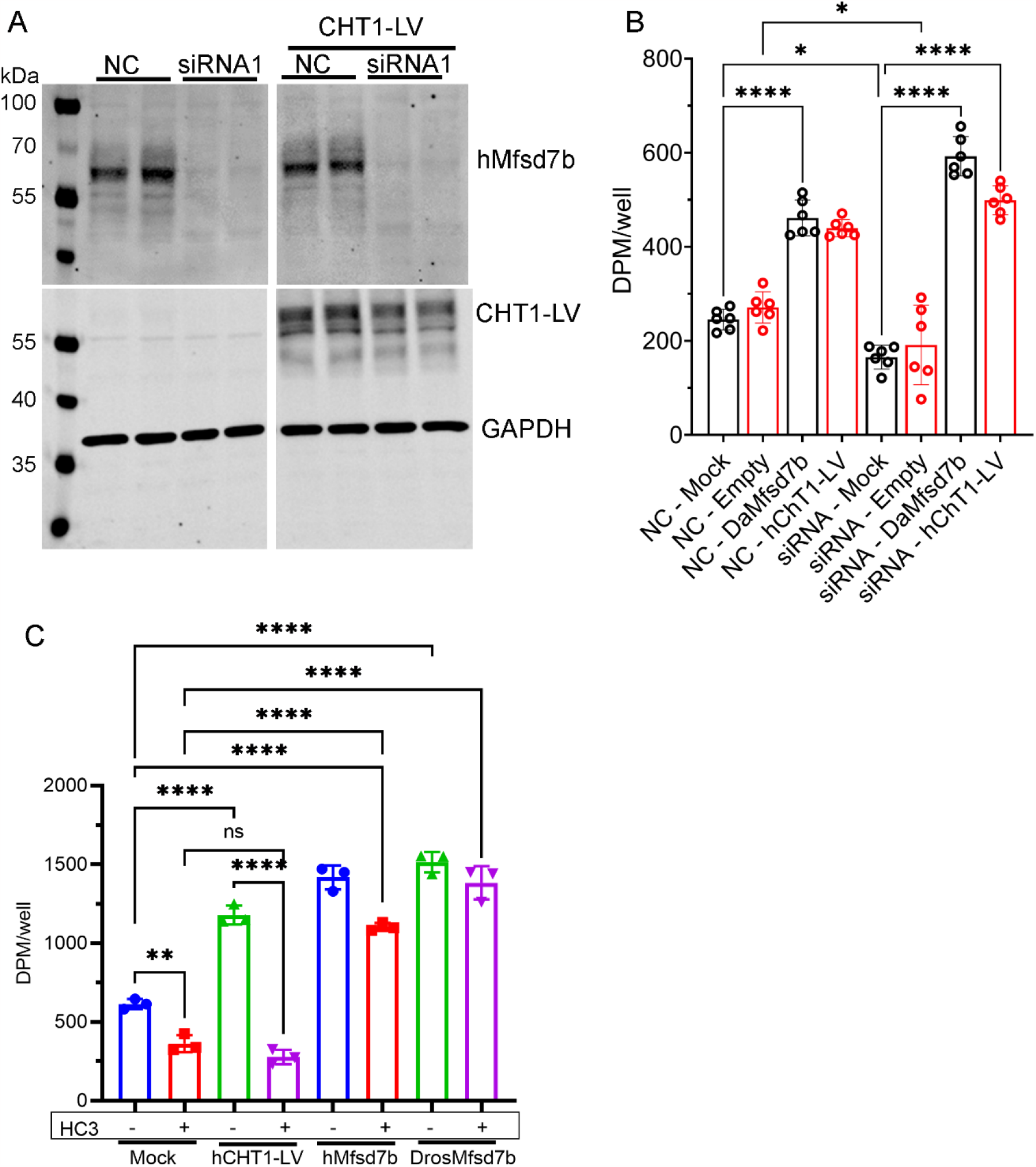
Mfsd7b functionally rescues for the reduced choline transport in HEK293 cells. **A-B**, Expression of zebrafish Mfsd7b or CHT1-LV rescued the reduced choline transport in Mfsd7b knockdown HEK293 cells. NC, negative controls; empty, pcDNA3.1 plasmid; mock, lipofectamine2000. **C**, Expression of human or fly Mfsd7b rescued the reduced choline transport via inhibition of CHT1, neuronal choline transporter in HEK293 cells. HEK293 cells were transfected with plasmids for human CHT1-LV dominant mutant (increased choline import activity), hMfsd7b, or drosophila Mfsd7b. The cells were assayed for choline import in the presence or absence of HC-3 to inhibit CHT1. Expression of hMfsd7b or drosophila Mfsd7b rescued the reduction of choline transport via CHT1 in HEK293 cells. Data are mean and SD. ****P<0.0001, **P<0.01, *P<0.05, ns, not significant. One-way ANOVA.

**Supplemental Figure 4.**
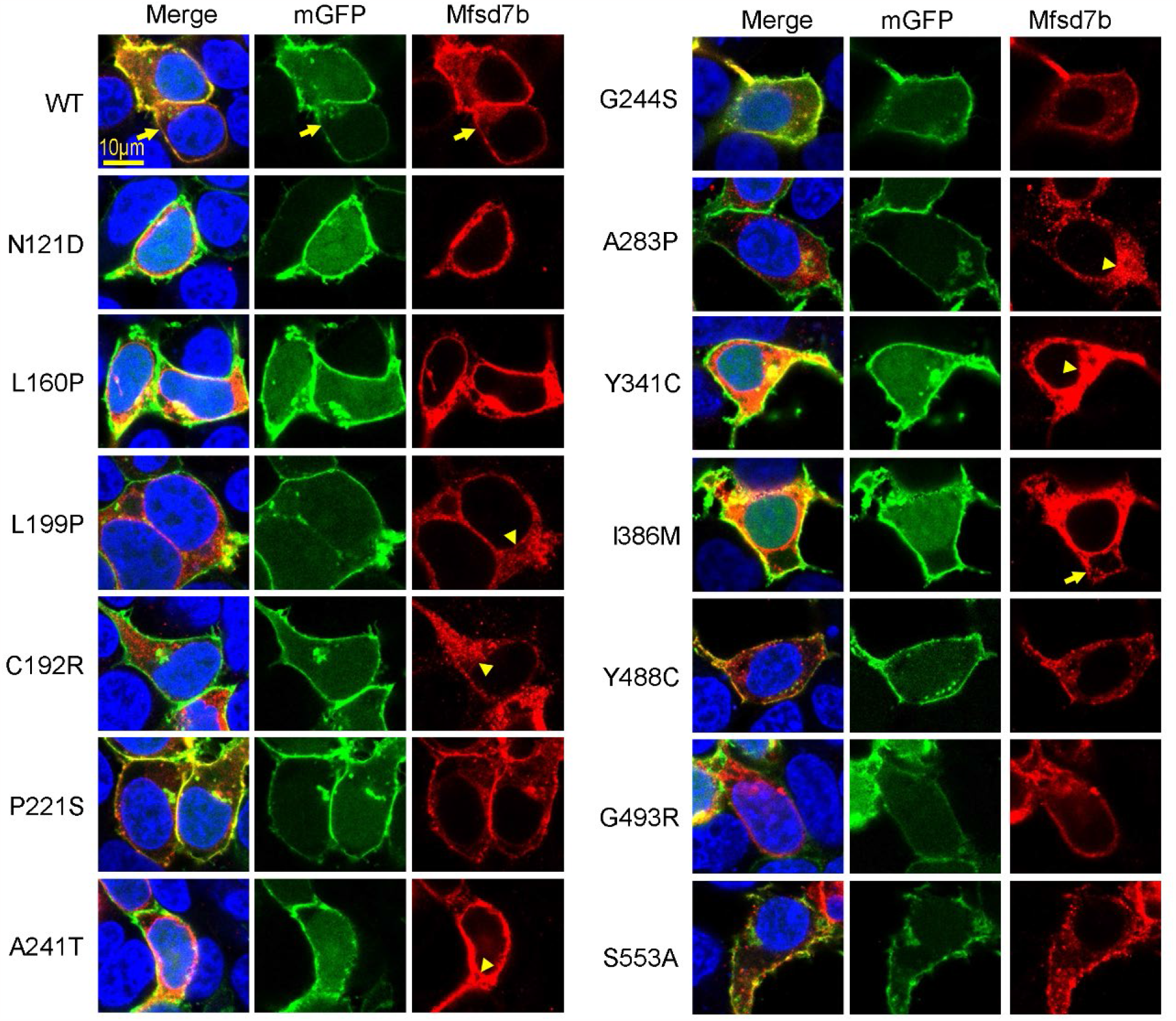
Immunofluorescence staining of hMFSD7b in HEK cells. HEK293 cells were overexpressed with hMFSD7b and stained with anti-hMFSD7b antibodies (red) and DAPI (blue). Membrane-localized GFP was also expressed to mark the plasma membrane. Localization of wildtype Mfsd7b (WT, arrows) and several Mfsd7b mutants appears to be in the plasma membrane, except for C192R, L199P, A241T, A283P, and Y341C mutants (arrowheads).

**Supplemental Table 1.**
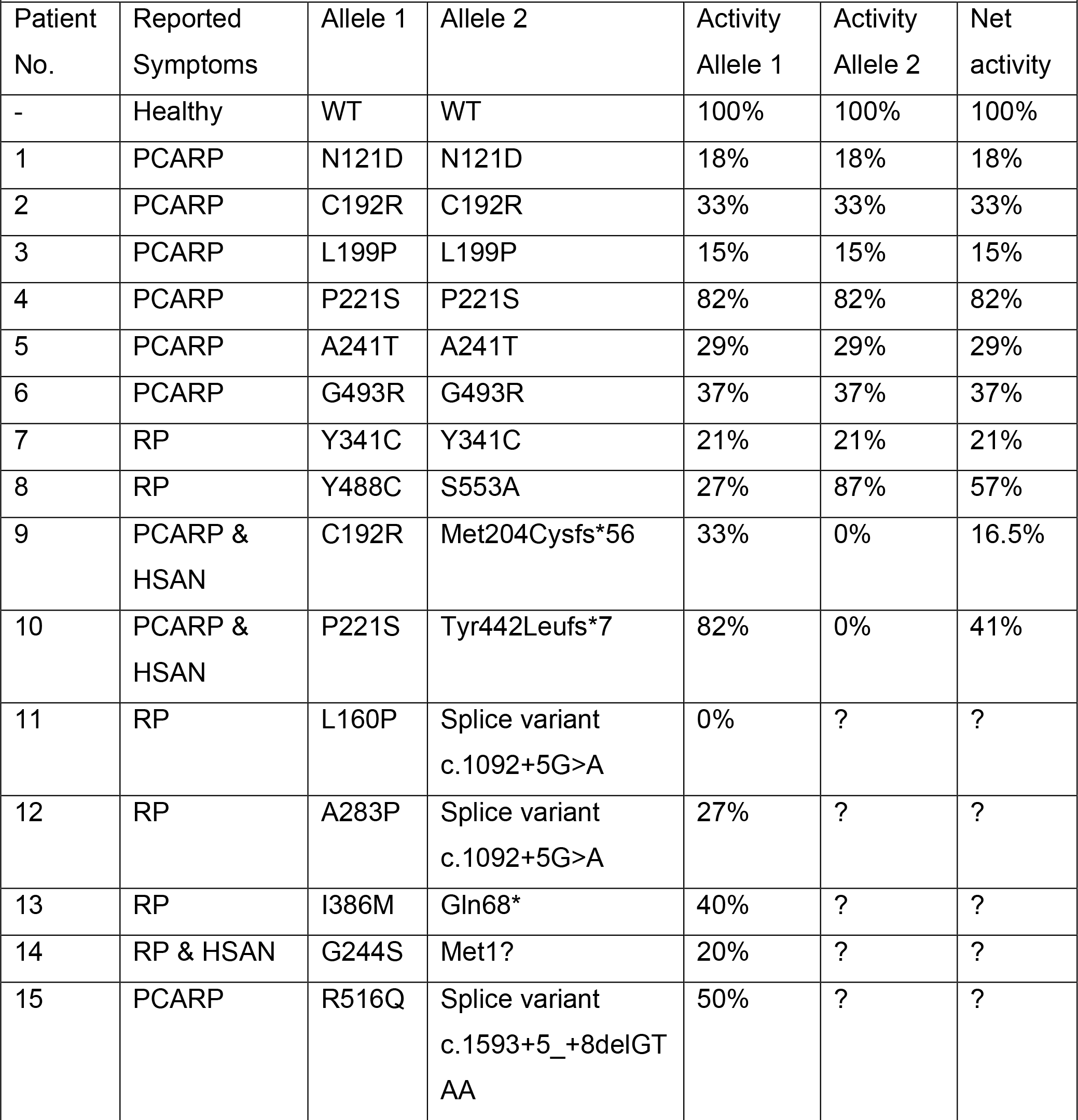
Summary of net choline import activity of MFSD7B in patients with PCARP, RP and HSAN. Net activity was calculated by taking the average choline import activity of the patients two reported MFSD7B allelic variants where possible.

| Patient No. | Reported Symptoms | Allele 1 | Allele 2 | Activity Allele 1 | Activity Allele 2 | Net activity |
| --- | --- | --- | --- | --- | --- | --- |
| - | Healthy | WT | WT | 100% | 100% | 100% |
| 1 | PCARP | N121D | N121D | 18% | 18% | 18% |
| 2 | PCARP | C192R | C192R | 33% | 33% | 33% |
| 3 | PCARP | L199P | L199P | 15% | 15% | 15% |
| 4 | PCARP | P221S | P221S | 82% | 82% | 82% |
| 5 | PCARP | A241T | A241T | 29% | 29% | 29% |
| 6 | PCARP | G493R | G493R | 37% | 37% | 37% |
| 7 | RP | Y341C | Y341C | 21% | 21% | 21% |
| 8 | RP | Y488C | S553A | 27% | 87% | 57% |
| 9 | PCARP & HSAN | C192R | Met204Cysfs*56 | 33% | 0% | 16.5% |
| 10 | PCARP & HSAN | P221S | Tyr442Leufs*7 | 82% | 0% | 41% |
| 11 | RP | L160P | Splice variant<br>c.1092+5G>A | 0% | ? | ? |
| 12 | RP | A283P | Splice variant<br>c.1092+5G>A | 27% | ? | ? |
| 13 | RP | I386M | Gln68* | 40% | ? | ? |
| 14 | RP & HSAN | G244S | Met1? | 20% | ? | ? |
| 15 | PCARP | R516Q | Splice variant<br>c.1593+5_+8delGT<br>AA | 50% | ? | ? |

## References

1. Quigley, J. G., Burns, C. C., Anderson, M. M., Lynch, E. D., Sabo, K. M., Overbaugh, J., and Abkowitz, J. L. (2000) Cloning of the cellular receptor for feline leukemia virus subgroup C (FeLV-C), a retrovirus that induces red cell aplasia. Blood 95, 1093–1099

2. Quigley, J. G., Yang, Z., Worthington, M. T., Phillips, J. D., Sabo, K. M., Sabath, D. E., Berg, C. L., Sassa, S., Wood, B. L., and Abkowitz, J. L. (2004) Identification of a human heme exporter that is essential for erythropoiesis. Cell 118, 757–766

3. Rajadhyaksha, A. M., Elemento, O., Puffenberger, E. G., Schierberl, K. C., Xiang, J. Z., Putorti, M. L., Berciano, J., Poulin, C., Brais, B., Michaelides, M., Weleber, R. G., and Higgins, J. J. (2010) Mutations in FLVCR1 cause posterior column ataxia and retinitis pigmentosa. Am J Hum Genet 87, 643–654

4. Ishiura, H., Fukuda, Y., Mitsui, J., Nakahara, Y., Ahsan, B., Takahashi, Y., Ichikawa, Y., Goto, J., Sakai, T., and Tsuji, S. (2011) Posterior column ataxia with retinitis pigmentosa in a Japanese family with a novel mutation in FLVCR1. Neurogenetics 12, 117–121

5. Yanatori, I., Yasui, Y., Miura, K., and Kishi, F. (2012) Mutations of FLVCR1 in posterior column ataxia and retinitis pigmentosa result in the loss of heme export activity. Blood Cells Mol Dis 49, 60–66

6. Chiabrando, D., Castori, M., di Rocco, M., Ungelenk, M., Giesselmann, S., Di Capua, M., Madeo, A., Grammatico, P., Bartsch, S., Hubner, C. A., Altruda, F., Silengo, L., Tolosano, E., and Kurth, (2016) Mutations in the Heme Exporter FLVCR1 Cause Sensory Neurodegeneration with Loss of Pain Perception. PLoS Genet 12, e1006461

7. Chiabrando, D., Marro, S., Mercurio, S., Giorgi, C., Petrillo, S., Vinchi, F., Fiorito, V., Fagoonee, S., Camporeale, A., Turco, E., Merlo, G. R., Silengo, L., Altruda, F., Pinton, P., and Tolosano, E. (2012) The mitochondrial heme exporter FLVCR1b mediates erythroid differentiation. J Clin Invest 122, 4569–4579

8. Keel, S. B., Doty, R. T., Yang, Z., Quigley, J. G., Chen, J., Knoblaugh, S., Kingsley, P. D., De Domenico, I., Vaughn, M. B., Kaplan, J., Palis, J., and Abkowitz, J. L. (2008) A heme export protein is required for red blood cell differentiation and iron homeostasis. Science 319, 825–828

9. Byon, J. C., Chen, J., Doty, R. T., and Abkowitz, J. L. (2013) FLVCR is necessary for erythroid maturation, may contribute to platelet maturation, but is dispensable for normal hematopoietic stem cell function. Blood 122, 2903–2910

10. Doty, R. T., Phelps, S. R., Shadle, C., Sanchez-Bonilla, M., Keel, S. B., and Abkowitz, J. L. (2015) Coordinate expression of heme and globin is essential for effective erythropoiesis. J Clin Invest 125, 4681–4691

11. Mercurio, S., Petrillo, S., Chiabrando, D., Bassi, Z. I., Gays, D., Camporeale, A., Vacaru, A., Miniscalco, B., Valperga, G., Silengo, L., Altruda, F., Baron, M. H., Santoro, M. M., and Tolosano, E. (2015) The heme exporter Flvcr1 regulates expansion and differentiation of committed erythroid progenitors by controlling intracellular heme accumulation. Haematologica 100, 720–729

12. Petrillo, S., Chiabrando, D., Genova, T., Fiorito, V., Ingoglia, G., Vinchi, F., Mussano, F., Carossa, S., Silengo, L., Altruda, F., Merlo, G. R., Munaron, L., and Tolosano, E. (2018) Heme accumulation in endothelial cells impairs angiogenesis by triggering paraptosis. Cell Death Differ 25, 573–588

13. Vinchi, F., Ingoglia, G., Chiabrando, D., Mercurio, S., Turco, E., Silengo, L., Altruda, F., and Tolosano, E. (2014) Heme exporter FLVCR1a regulates heme synthesis and degradation and controls activity of cytochromes P450. Gastroenterology 146, 1325–1338

14. Fiorito, V., Forni, M., Silengo, L., Altruda, F., and Tolosano, E. (2015) Crucial Role of FLVCR1a in the Maintenance of Intestinal Heme Homeostasis. Antioxid Redox Signal 23, 1410–1423

15. Tsuchiya, M., Tachibana, N., Nagao, K., Tamura, T., and Hamachi, I. (2023) Organelle-selective click labeling coupled with flow cytometry allows pooled CRISPR screening of genes involved in phosphatidylcholine metabolism. Cell Metab 35, 1072–1083 e1079

16. Kenny, T. C., Khan, A., Son, Y., Yue, L., Heissel, S., Sharma, A., Pasolli, H. A., Liu, Y., Gamazon, E. R., Alwaseem, H., Hite, R. K., and Birsoy, K. (2023) Integrative genetic analysis identifies FLVCR1 as a plasma-membrane choline transporter in mammals. Cell Metab 35, 1057–1071 e1012

17. An, X., Schulz, V. P., Li, J., Wu, K., Liu, J., Xue, F., Hu, J., Mohandas, N., and Gallagher, P. G. (2014) Global transcriptome analyses of human and murine terminal erythroid differentiation. Blood 123, 3466–3477

18. Fiorito, V., Allocco, A. L., Petrillo, S., Gazzano, E., Torretta, S., Marchi, S., Destefanis, F., Pacelli, C., Audrito, V., Provero, P., Medico, E., Chiabrando, D., Porporato, P. E., Cancelliere, C., Bardelli, A., Trusolino, L., Capitanio, N., Deaglio, S., Altruda, F., Pinton, P., Cardaci, S., Riganti, C., and Tolosano, E. (2021) The heme synthesis-export system regulates the tricarboxylic acid cycle flux and oxidative phosphorylation. Cell Rep 35, 109252

19. Yachie, A., Niida, Y., Wada, T., Igarashi, N., Kaneda, H., Toma, T., Ohta, K., Kasahara, Y., and Koizumi, S. (1999) Oxidative stress causes enhanced endothelial cell injury in human heme oxygenase-1 deficiency. J Clin Invest 103, 129–135

20. Radhakrishnan, N., Yadav, S. P., Sachdeva, A., Pruthi, P. K., Sawhney, S., Piplani, T., Wada, T., and Yachie, A. (2011) Human heme oxygenase-1 deficiency presenting with hemolysis, nephritis, and asplenia. J Pediatr Hematol Oncol 33, 74–78

21. Chau, A. S., Cole, B. L., Debley, J. S., Nanda, K., Rosen, A. B. I., Bamshad, M. J., Nickerson, D. A., Torgerson, T. R., and Allenspach, E. J. (2020) Heme oxygenase-1 deficiency presenting with interstitial lung disease and hemophagocytic flares. Pediatr Rheumatol Online J 18, 80

22. Poss, K. D., and Tonegawa, S. (1997) Heme oxygenase 1 is required for mammalian iron reutilization. Proc Natl Acad Sci U S A 94, 10919–10924

23. Tavasoli, M., Lahire, S., Reid, T., Brodovsky, M., and McMaster, C. R. (2020) Genetic diseases of the Kennedy pathways for membrane synthesis. J Biol Chem 295, 17877–17886

24. Choudhary, P., Armstrong, E. J., Jorgensen, C. C., Piotrowski, M., Barthmes, M., Torella, R., Johnston, S. E., Maruyama, Y., Janiszewski, J. S., Storer, R. I., Skerratt, S. E., and Benn, C. L. (2017) Discovery of Compounds that Positively Modulate the High Affinity Choline Transporter. Front Mol Neurosci 10, 40

